# Tobacco, but not nicotine and flavor-less electronic cigarettes, induces ACE2 and immune dysregulation

**DOI:** 10.1101/2020.07.13.198630

**Authors:** Abby C. Lee, Jaideep Chakladar, Wei Tse Li, Chengyu Chen, Eric Y. Chang, Jessica Wang-Rodriguez, Weg M. Ongkeko

**Author notes:** Authors contributed equally (Abby C. Lee, Jaideep Chakladar, Wei Tse Li).

## Abstract

COVID-19, caused by the virus SARS-CoV-2, has infected millions worldwide. This pandemic overlaps with the ongoing epidemics of cigarette smoking and electronic cigarette (e-cig) vaping, with over 1 billion smokers and vapers worldwide. However, there is scarce data relating COVID-19 risks and outcome with cigarette or e-cig use. In this study, we mined 3 independent RNA expression datasets from smokers and vapers to understand the potential relationship between vaping/smoking and the dysregulation of key genes and pathways related to COVID-19. We found that smoking, but not vaping, upregulates ACE2, the cellular receptor that SARS-CoV-2 requires for infection. Both smoking and use of nicotine and flavor-containing e-cig led to upregulations of pro-inflammatory cytokine production and expression of genes related to inflammasomes. Vaping flavor-less and nicotine-less e-cig, however, did not lead to significant cytokine dysregulation and inflammasome activation. Release of inflammasome products, such as IL-1B, and cytokine storms are hallmarks of COVID-19 infection, especially in severe cases. Therefore, our findings demonstrated that smoking or vaping, specifically use of flavored or nicotine-containing e-cigs, may critically exacerbate COVID-19-related inflammation or increase susceptibility to the disease. Further scientific and public health investigations should be undertaken to address these concerning links between COVID-19 and e-cig/smoking.

## 1. Introduction

The current COVID-19 pandemic, caused by the virus SARS-CoV-2, has killed over 130,000 Americans in less than 4 months. The majority are over 65 years old, but many younger patients who have underlying medical conditions required admission to the intensive care unit. Current data indicate that patients who have cardiovascular and chronic respiratory conditions, including those caused by tobacco use, are at higher risk of developing severe COVID-19 symptoms and have significantly increased fatality [1]. There are increasing numbers of reports that smokers have worse clinical outcomes when infected with SARS-CoV-2 [2]. Information from China show that people who have cardiovascular and respiratory conditions caused by tobacco use are at higher risk of developing severe COVID-19 symptoms [2,3]. Furthermore, tobacco use is the most important risk-factor for chronic obstructive pulmonary disease (COPD) [4]. Finally, aside from lung damage, it is believed that smokers may be more vulnerable to SARS-CoV-2 because of an altered immune response [5]. However, many studies reported otherwise. In a preprint, Farsalinos et. al. concluded that smoking is not a risk factor for hospitalization due to COVID-19 [6]. One meta-analysis reported no association between smoking and severity of COVID-19 [7]. A French study even found that nicotine may protect against COVID-19 infection, prompting interests from the French Health Minister in studying whether nicotine patches could be used for COVID-19 treatment and prevention [8,9]. Unfortunately, all reports regarding tobacco and COVID-19 are from epidemiological data and only a few studies reported data. The lack of well-controlled laboratory experiments renders it extremely difficult to determine whether tobacco truly affects COVID-19 outcome and the mechanistic link between them. Furthermore, no peered-reviewed study has reported definitive data demonstrating or rejecting the hypothesis that COVID-19 incidence can be increased by tobacco smoking.

Electronic cigarettes (e-cigs) have dramatically increased in popularity in recent years, especially among the youth [10], so considerable attention has been directed to any potential link between e-cig and COVID-19 in recent months. E-cig has been shown to induce inflammation of human airways and may also increase susceptibility to pneumonia by increasing pneumococcal adherence to airway cells [11,12]. Notable damage to the lung epithelium could also be done by e-cig, resulting in increased airway hyperactivity and mucine production [12]. The detrimental effects of e-cig thus fueled speculation that e-cig may increase COVID-19 severity or incidence. However, virtually no study, both on the basic science level and epidemiological level, has linked COVID-19 to e-cig. Complicating research on e-cig’s effects on COVID-19 is the fact that e-cig can be vaped with or without nicotine and with or without flavorings. Flavors and nicotine in e-cig are frequently associated with more inflammation, epithelial barrier dysregulation, oxidative stress, and DNA damage than e-cig with only basic components [12,13].

A possible molecular mechanism for tobacco or e-cig to increase COVID-19 susceptibility is the upregulation of ACE2. For SARS-CoV-2, host recognition is carried out by the spike protein on the surface of the viral envelope. The spike protein binds the ACE2 receptor protein in human cells. After this, the spike protein can be cleaved by the serine protease TMPRSS2 [14]. SARS-CoV uses a similar mechanism for host recognition, after which, fusion of the viral envelope with membranes in the host cell will allow the virus entry into the cell [15]. This process is vital for the entry of SARS-CoV-2 into human host cells, and therefore plays an integral role in COVID-19 infection and disease progression. Cigarette smoke was found in some studies to increase ACE2 expression in lungs in mammals [16–18]. No study has reported on ACE2 expression’s relationship with E-cig use.

Another possible mechanism linking tobacco or cigarette smoking to COVID-19 is their propensity to increase lung inflammation. COVID-19 has a strong immunological component, and poor outcomes have recently been associated with cytokine storms and a hyperinflammatory immune system [19,20]. Tobacco is known to induce extensive immune dysregulation in smokers, and its influence has been found to lead to airway inflammation[21]. Although cigarette smoke has been associated with increased susceptibility to COVID-19[16], the effect of tobacco on the immune response to SARS-CoV-2 has not been studied. E-cig is also known to induce significant inflammation in the lungs, but the dysregulated immune landscape is different from that of tobacco[22,23]. Therefore, it is plausible that inflammation induced by tobacco or e-cig could exacerbate inflammation caused by SARS-CoV-2 infection, potentially triggering a lethal cytokine storm.

In this study, we directly investigated the two main hypotheses that might link COVID-19 to e-cig vaping and tobacco smoking, ACE2 upregulation and inflammation, by reanalysis of 3 independent datasets with gene expression from e-cig vapers and tobacco smokers. We examined whether e-cig or tobacco upregulate ACE2, dysregulate immune cell population levels, affect cytokine levels, and regulate the activation of inflammasomes. Given that nicotine or flavorings present in the e-cig mixture are implicated in promoting lung damage, we examined one e-cig dataset where all participants only vaped e-cig with neither nicotine nor flavors and another e-cig dataset where participants only vaped e-cig with nicotine and were free to choose different flavors.

## 2. Results

### 2.1. Correlation between smoking/vaping and ACE2 expression in bronchial epithelial cells

To observe the relationship between smoking e-cigarettes and ACE2 expression, we used the following datasets: GSE138326 and GSE112073 [24,25]. The GSE138326 dataset, from Song et. al., details gene expression in the bronchial epithelial cells of patients who smoked flavor-less and nicotine-less e-cig vs. those who did not. The GSE112073 dataset, from Corbett et. al., details gene expression in bronchial cells of patients who smoked nicotine-containing e-cig of any flavor vs. those who smoked no e-cig. All patients in this dataset were former smokers. The Kruskal-Wallis analysis test was applied to determine differential expression of ACE2 between e-cig users and non-e-cig users for both of these datasets (p<0.05). In addition, whole transcriptome RNA sequencing data for normal bronchus and lung tissue samples of 49 lung squamous cell carcinoma (LUSC) patients from The Cancer Genome Atlas (TCGA) were downloaded. These patients are either current or former smokers. Former smokers were categorized into those who had quit less than 15 years from the date of sample collection and those who had quit more than 15 years. Differential expression analysis was conducted to compare gene expression in current smokers and former smokers (Kruskal-Wallis, p<0.05).

We found that smoking tobacco is associated with increased ACE2 expression in current smokers (p=0.0281). In addition, the difference in ACE2 expression between current smokers and former smokers who had quit for more than 15 years is more significant than that of current smokers and former smokers who had quit for less than 15 years (Fig 1A). This is not surprising as we expect that a longer duration since quitting smoking would be associated with a greater difference in ACE2 expression between current and former smokers. On the other hand, smoking e-cigarettes does not lead to increased ACE2 expression in either study (Fig 1B). These results suggest that tobacco smokers may be more vulnerable to SARS-CoV-2 infection than e-cigarette smokers if infection susceptibility is based upon ACE2 abundance.

**Figure 1:**
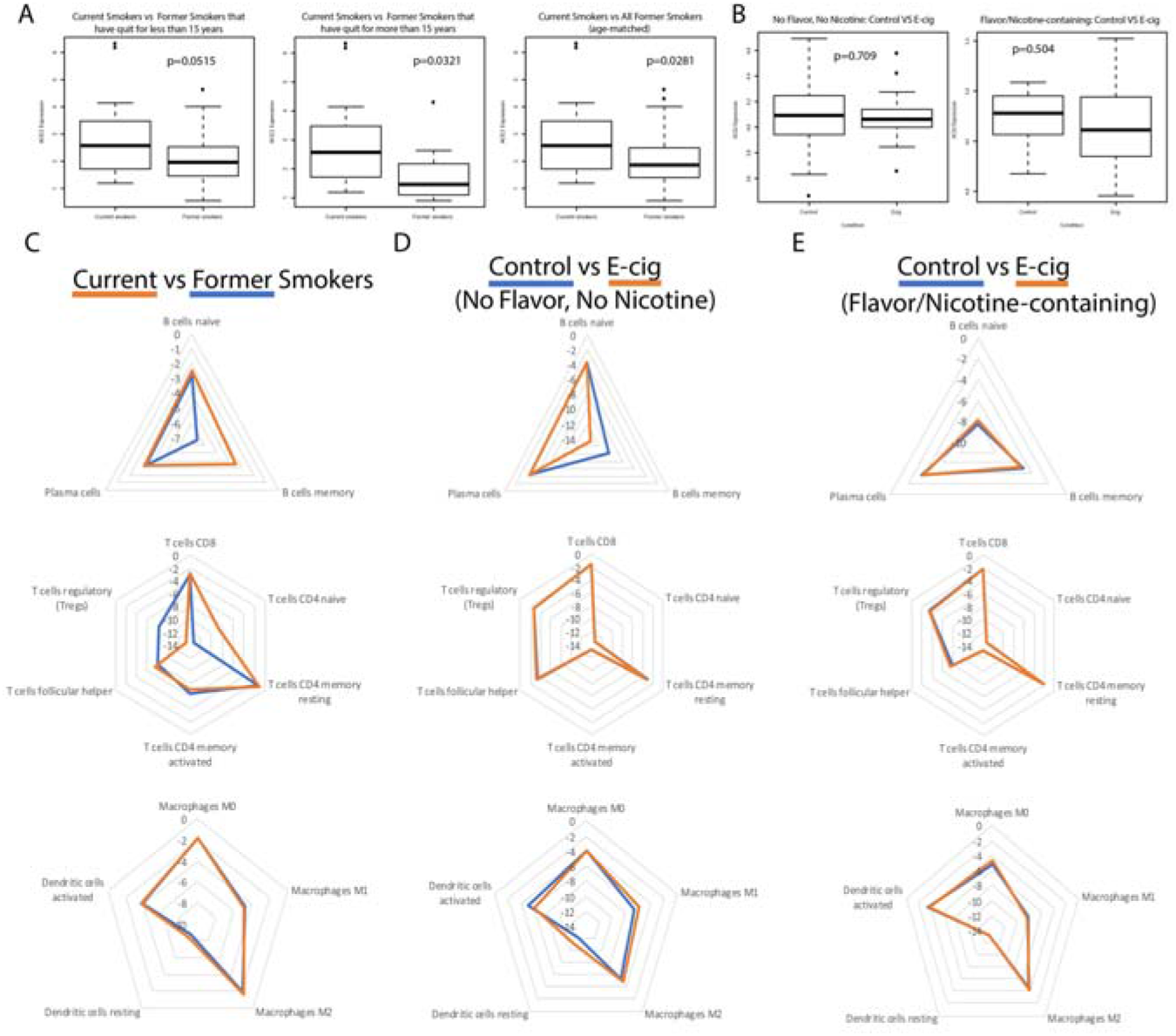
ACE2 and general immune dysregulation analysis in tobacco and e-cig users. (A) Boxplots of ACE2 expression in current and former smokers, including those that have quit for less than and more than 15 years. (B) Boxplots of ACE2 expression in control and e-cigarette users, including patients who have never smoked tobacco and those who are former tobacco smokers. (C-E) Radar plots comparing immune cell infiltration across smokers, e-cigarette users vaping nicotine-less, flavor-less e-cig, and vapers who used e-cigs containing nicotine and flavors. Greater immune cell infiltration is indicated by a value closer to zero and farther away from the center of the radar plot.

### 2.2. Correlation between smoking/vaping status and immune infiltration

Using the software tool CIBERSORTx [26], we deconvoluted the infiltrating lymphocyte population from bulk-tissue gene expression data (p<0.05). We also found that smoking tobacco is associated with increased immune cell infiltration (Fig 1C-E). Current tobacco smokers experienced an upregulation of memory B-cell and naive CD4 T-cell expression and downregulation of regulatory T-cell (Treg) expression, compared to former smokers (Figure 1C). Elevated levels of CD4 T-cells in current tobacco smokers could lead to higher levels of effector cytokines that contribute to lung inflammation, making tobacco smokers more susceptible to a more aggressive immune response and a possible cytokine storm. In fact, it was reported that CD4+ T-cells contribute effector inflammation cytokines after acute influenza infections in mice, even after no virus is detectable in blood [27]. Meanwhile, the downregulation of Tregs may deprive the immune system of inhibitory signals that can prevent cytokine storms from occurring [28]. Memory B cells upregulation is not known to be involved in a cytokine storm, but their expansion may be indicative of past infections and increased susceptibility to infection in smokers. Our data is consistent with existing literature, which reports that current smokers have higher percentages of memory B cells than former smokers [29]. On the other hand, e-cig use is not significantly associated with increased immune infiltration in either datasets (Fig 1D-E).

### 2.3. Correlation between smoking/vaping status and cytokine level dysregulation

We next examined whether cytokine levels are dysregulated in smokers or e-cig users. We found that current tobacco smokers display significant dysregulation of 18 cytokine genes, including 12 proinflammatory and 6 anti-inflammatory cytokines, compared to former tobacco smokers (Fig 2A,D,E). Of the proinflammatory cytokines, the following are of particular interest: CCL20, which has been associated with SARS-CoV, and IL-1B, CXCL1, CXCL2 and CXCL8, which are associated with COVID-19. CCL20 was highly expressed in healthy Peripheral Blood Mononuclear Cells (PBMCs) infected with SARS-CoV [30]. Given the similarities in the pathophysiology of SARS-CoV and COVID-19 infection, more research must be done to determine if CCL20 is upregulated in COVID-19 patients as well, thus contributing to a potential cytokine storm. COVID-19 infection of the upper and lower respiratory tract has been shown to release proinflammatory cytokines including IL-1B [31], and is also associated with increased levels of CXCL8, CXCL1, and CXCL2 [32], which are known to play roles in neutrophil recruitment [33,34]. CXCL8 has also been found to be positively correlated with the number of neutrophils recovered in ARDS patients [34]. Such excessive neutrophil recruitment during ARDS is associated with poorer clinical outcomes and greater disease severity [35,36]. Therefore, the upregulation of these cytokines in tobacco smokers may make them more vulnerable to excessive neutrophil migration, a more severe inflammatory response, and leave them more susceptible to cytokine storms. Of the anti-inflammatory cytokines, CXCL17 is upregulated in COVID-19 patients as well [32], but more research must be done to investigate its role in COVID-19 pathogenesis.

**Figure 2:**
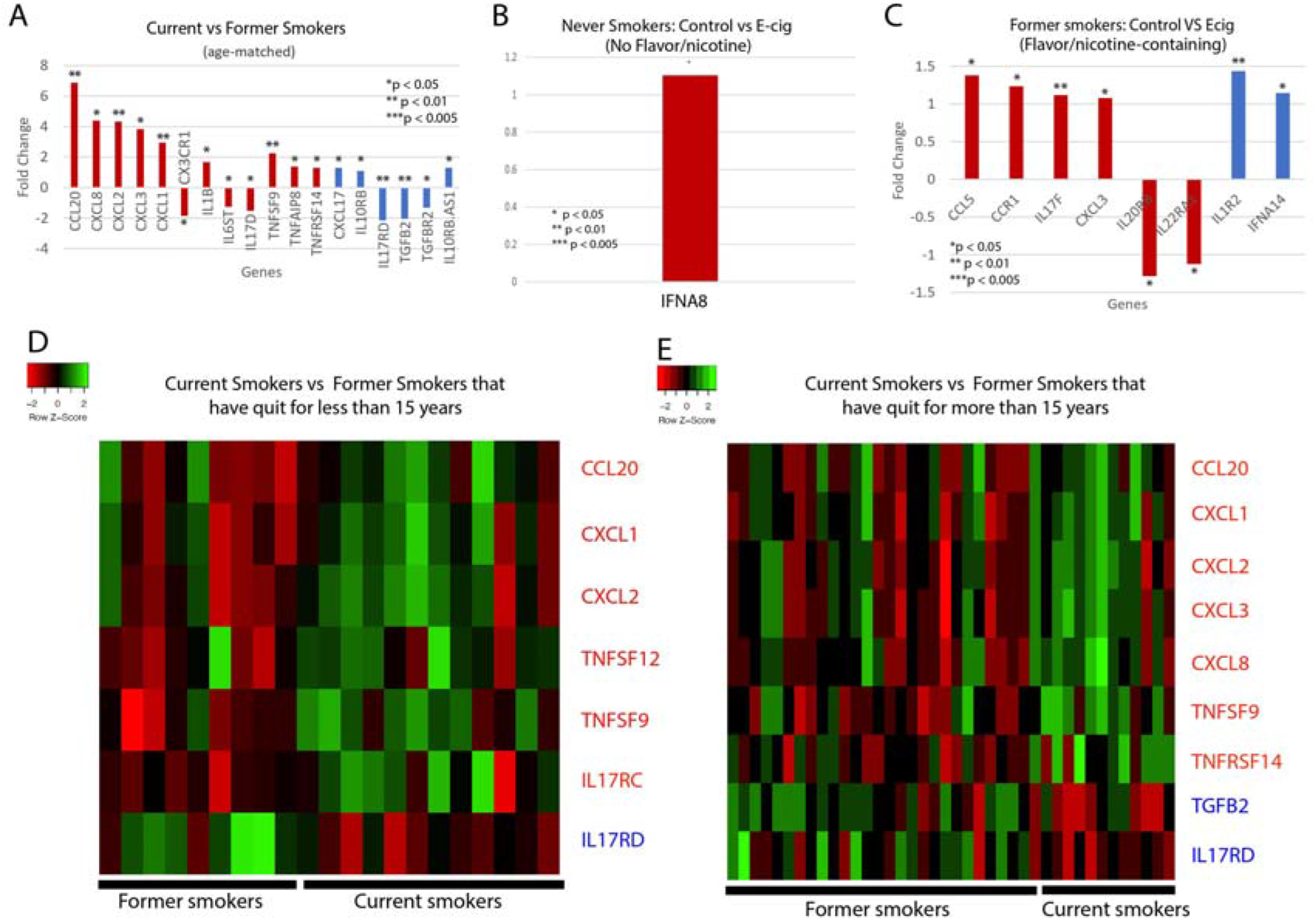
Cytokine-related gene expression. Bar plots of the fold changed of significantly dysregulated cytokine genes in (A) Current vs former smokers; and control vs e-cig users that have used e-cig with (B) no flavors or nicotine or (C) with nicotine and/or flavors. Heat maps illustrating differences in cytokine gene expression between current smoker and former smokers that have either quit for (D) less than 15 years or (E) more than 15 years. Heat maps are split into quadrants based on current/ former smokers and pro/anti-inflammatory cytokines. Red bar plots represent proinflammatory cytokines, and blue bar plots represent anti-inflammatory cytokines.

For the e-cig studies, we noticed that there are significant differences between vapers who used nicotine-less and flavor-less e-cig and vapers who used nicotine/flavor-containing e-cig. In nicotine-less and flavor-less e-cig users, we observed only one cytokine, IFNA8, being slightly dysregulated (Figure 2B). However, we observed the dysregulation of 6 pro-inflammatory cytokines, including CCL5, CCL3, and CXCL3, and 2 anti-inflammatory cytokines in those who used nicotine/flavor-containing e-cig (Figure 2C). CCL5 and CCL3 are both chemotactic cytokines that specifically attract T-cells and macrophages to the site of infection [37,38]. CCL5 is also critical to inhibiting macrophages from undergoing apoptosis after the activation of adaptive immunity, potentially leading to prolonged inflammation [38]. CXCL3 is a chemokine released by macrophages and may indicate increased airway inflammation [39]. In summary, our data suggest that use of nicotine or flavor-containing e-cig may lead to greater expression of inflammatory genes.

**Table 1:**
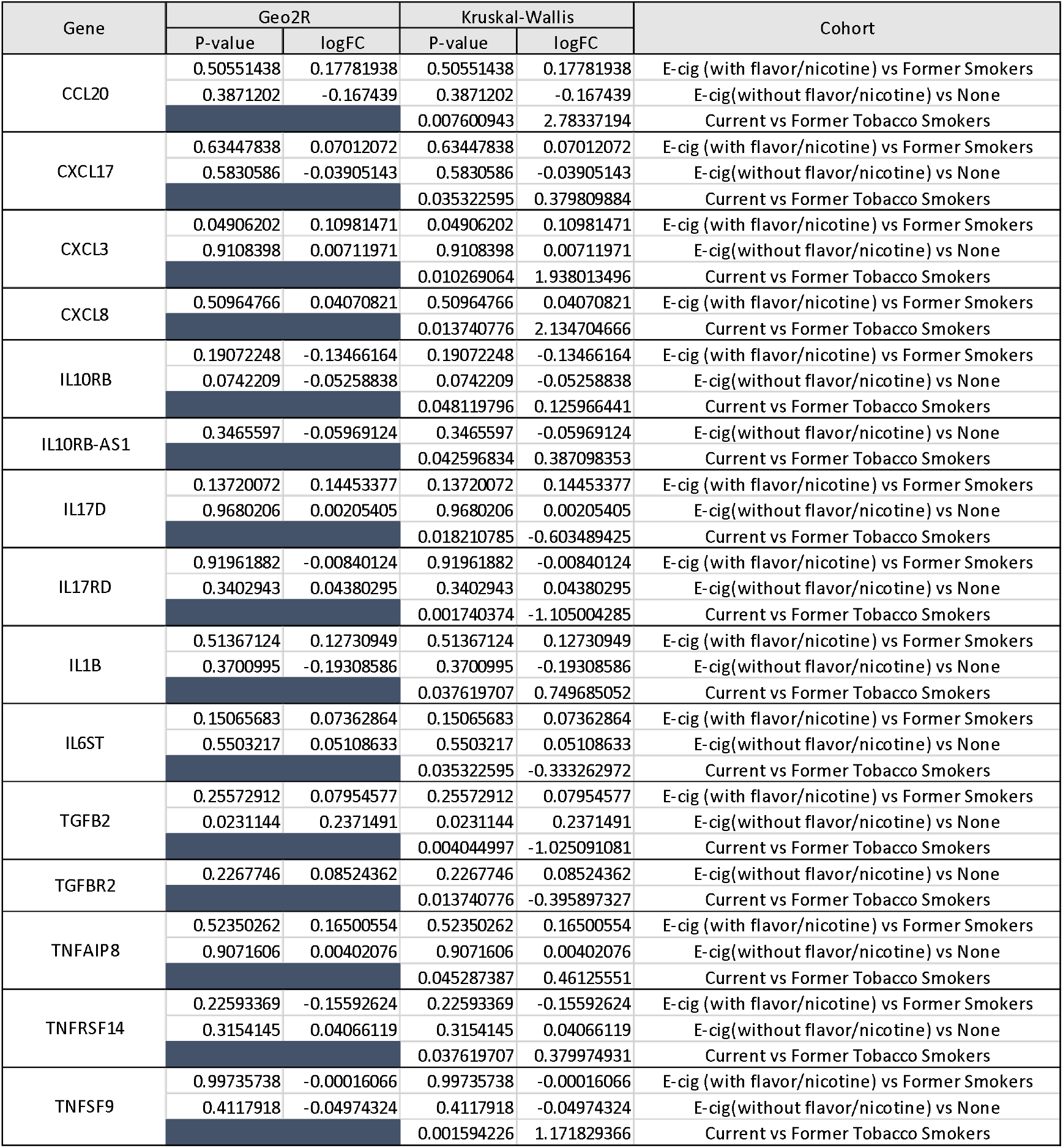
Significantly dysregulated Cytokine-related genes.

### 2.4. Investigation of inflammasome activation in e-cig and tobacco users

The upregulation of a significant number of inflammatory cytokines in smokers and nicotine/flavor-containing e-cig users and the association of smoking with IL-1B prompted us to examine inflammasome activation in smokers and e-cig users. Inflammasomes are large protein structures primarily located in macrophages that responds to inflammatory signals, namely molecular patterns from pathogens. When activated, inflammasomes cleave pro-IL-1B and pro-IL18 into IL-1B and IL18, allowing them to signal the presence of pathogens to nearby cells and initate the inflammatory response [40]. The discovery that IL-1B is upregulated by both smoking and COVID-19 suggests the possibility that inflammasomes are activated.

**Table 2:**
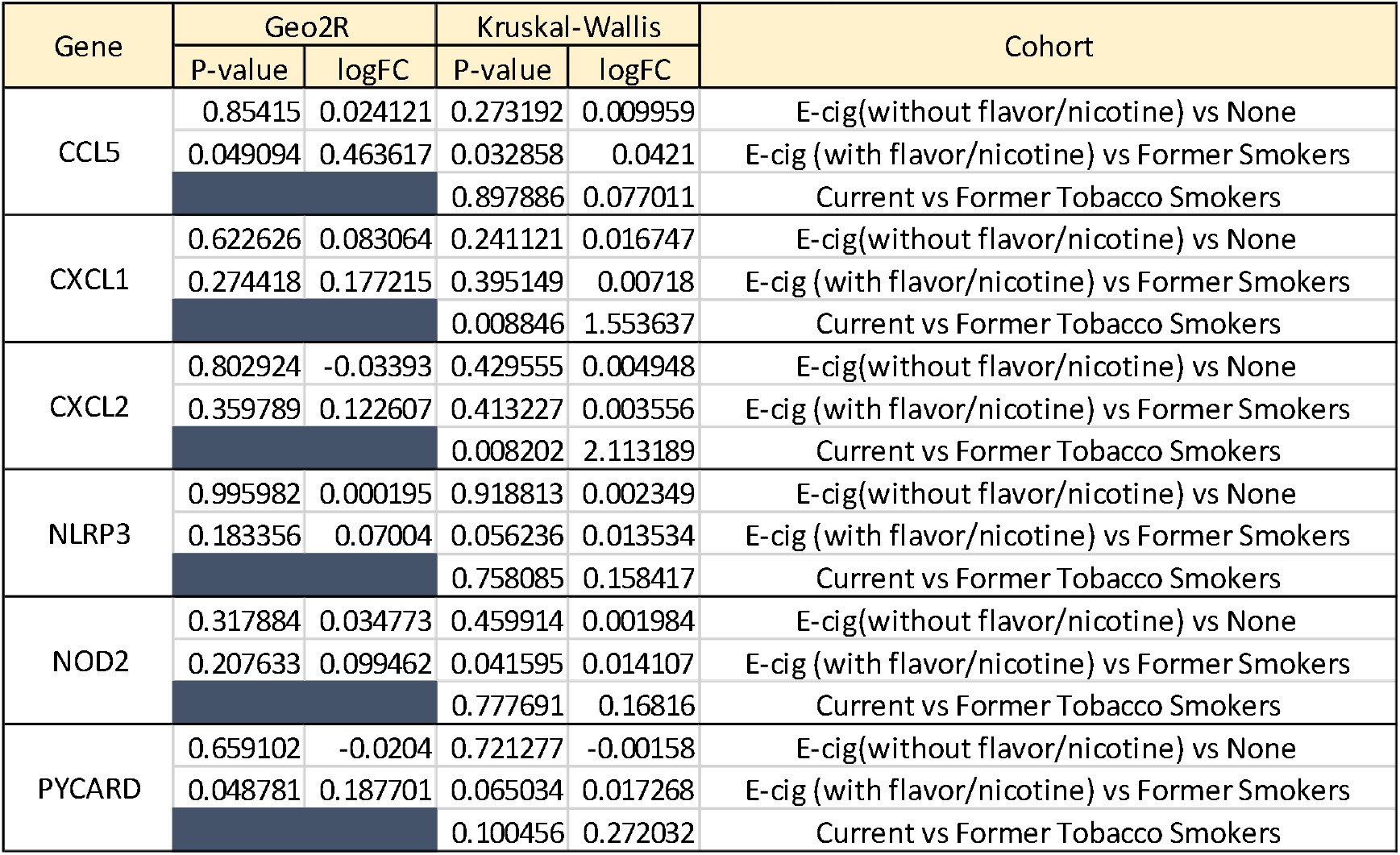
Significantly dysregulated Inflammasome-related gene expression

| Gene | Geo2R |  | Kruskal-Wallis |  | Cohort |
| --- | --- | --- | --- | --- | --- |
|  | P-value | logFC | P-value | logFC |  |
| CCL5 | 0.85415 | 0.024121 | 0.273192 | 0.009959 | E-cig(without flavor/nicotine) vs None |
|  | 0.049094 | 0.463617 | 0.032858 | 0.0421 | E-cig (with flavor/nicotine) vs Former Smokers |
|  |  |  | 0.897886 | 0.077011 | Current vs Former Tobacco Smokers |
| CXCL1 | 0.622626 | 0.083064 | 0.241121 | 0.016747 | E-cig(without flavor/nicotine) vs None |
|  | 0.274418 | 0.177215 | 0.395149 | 0.00718 | E-cig (with flavor/nicotine) vs Former Smokers |
|  |  |  | 0.008846 | 1.553637 | Current vs Former Tobacco Smokers |
| CXCL2 | 0.802924 | -0.03393 | 0.429555 | 0.004948 | E-cig(without flavor/nicotine) vs None |
|  | 0.359789 | 0.122607 | 0.413227 | 0.003556 | E-cig (with flavor/nicotine) vs Former Smokers |
|  |  |  | 0.008202 | 2.113189 | Current vs Former Tobacco Smokers |
| NLRP3 | 0.995982 | 0.000195 | 0.918813 | 0.002349 | E-cig(without flavor/nicotine) vs None |
|  | 0.183356 | 0.07004 | 0.056236 | 0.013534 | E-cig (with flavor/nicotine) vs Former Smokers |
|  |  |  | 0.758085 | 0.158417 | Current vs Former Tobacco Smokers |
| NOD2 | 0.317884 | 0.034773 | 0.459914 | 0.001984 | E-cig(without flavor/nicotine) vs None |
|  | 0.207633 | 0.099462 | 0.041595 | 0.014107 | E-cig (with flavor/nicotine) vs Former Smokers |
|  |  |  | 0.777691 | 0.16816 | Current vs Former Tobacco Smokers |
| PYCARD | 0.659102 | -0.0204 | 0.721277 | -0.00158 | E-cig(without flavor/nicotine) vs None |
|  | 0.048781 | 0.187701 | 0.065034 | 0.017268 | E-cig (with flavor/nicotine) vs Former Smokers |
|  |  |  | 0.100456 | 0.272032 | Current vs Former Tobacco Smokers |

We discovered that many key inflammasome genes and regulators besides IL-1B were upregulated in smokers vs. former smokers, including CXCL1 and CXCL2 (Figure 3). These genes are all upregulated in smokers vs. former smokers. Both CXCL1 and CXCL2 promote the activation of inflammasomes [41,42]. In the e-cig studies, we found that flavor-less and nicotine-less e-cig users showed no markers of inflammasome activation (Figure 3). However, users of nicotine/flavor-containing e-cig also exhibited highly significant upregulation of inflammasome-related genes, including NOD2, CCL5, and ASC (also known as PYCARD) (Figure 3). ASC is one of the 3 main components of inflammasomes. NOD2 can direcly bind to NLRP1 inflammasomes to upregulate IL-1B release [43]. CCL5 expression may be induced by inflammasome activation, as IL-1B has been shown to induce CCL5 expression in lower airway cells [44]. Our data thus suggest that smoking and nicotine or flavor-containing e-cig, but not nicotine/flavor-less e-cig, could upregulate inflammasomes and potentially exacerbate COVID-19-induced inflammation.

**Figure 3:**
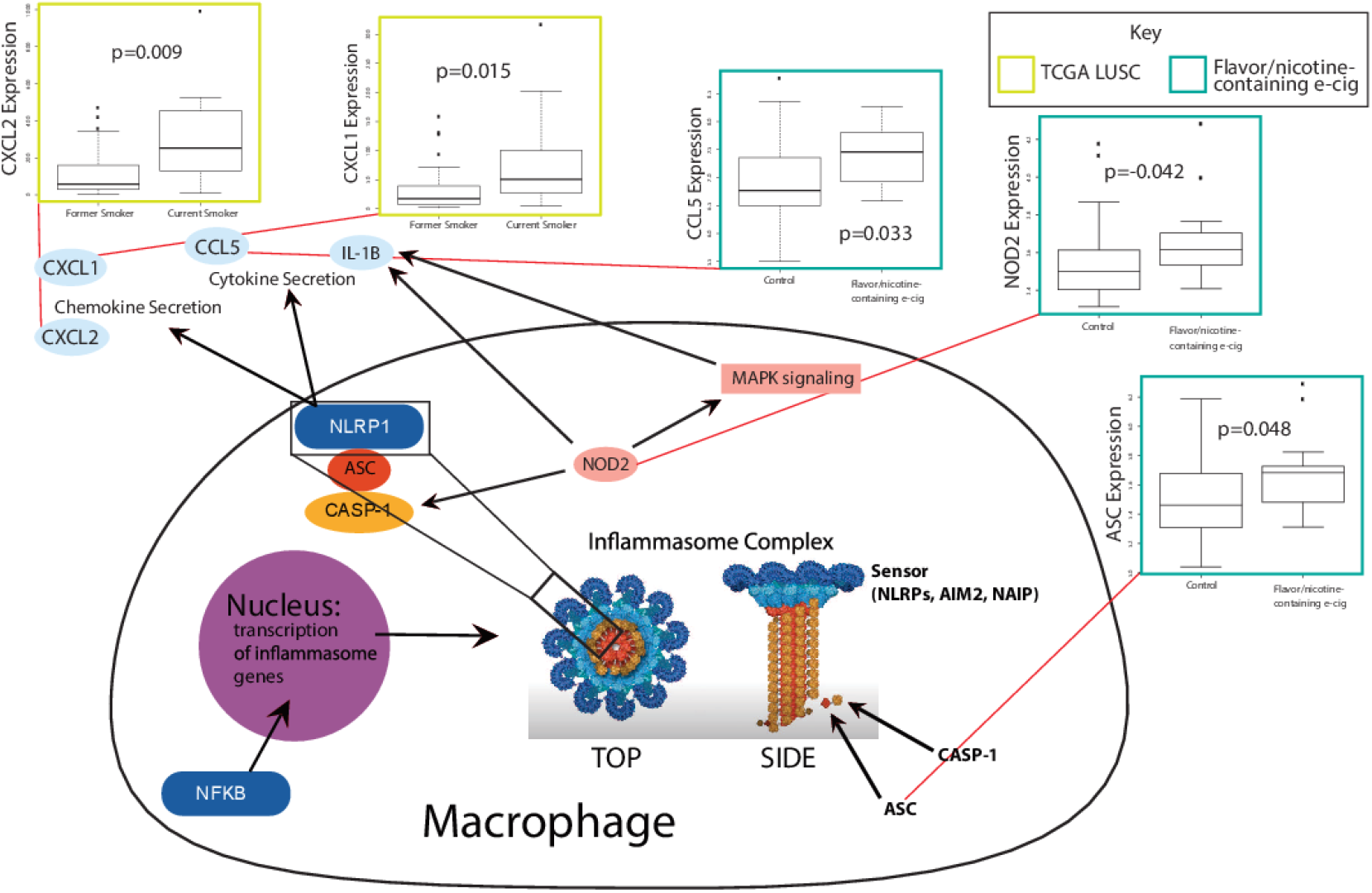
Boxplots of significant correlations of e-cig or tobacco use with upregulation of inflammasome genes (Kruskal Wallis, *p*<0.05). A schematic linking the genes upregulated with their function within inflammasome pathways is shown.

## 3. Discussion

The current COVID-19 pandemic intersects with the ongoing epidemic of tobacco smoking and e-cig vaping in the U.S. and across the globe. Tobacco usage has been declining gradually over the years. 42.3% of U.S. adults smoked cigarettes in 1965, but only 13.7 % do so in 2018, according to data from the National Health Interview Survey (NHIS). The smoking rate among young adults have also fallen similarly, with 7% of young adults smoking in 2018 [44]. Although these trends are encouraging, it does not mask the fact that millions of Americans are still active smokers. The many detrimental effects tobacco can have on the lungs and other organs have made tobacco usage one of the biggest public healthy crises the world faces. Accumulating evidence suggests that smokers are more susceptible to bacterial or viral infection and exhibit greater severity of these infections [45,46]. There have also been limited accounts of cigarette smoking increasing severity of COVID-19 infection [46]. The millions of smokers worldwide may be especially vulnerable to COVID-19.

However, more concerning than tobacco usage has been the usage of e-cig. Although not nearly as popular as tobacco, e-cig usage has been gaining traction at alarming rates. Critically, the usage of e-cig has been increasing dramatically among teenagers. According to CDC data, more than 1.3 million high school students started using e-cig in one year alone (2017-2018), which is a 78% increase in usage (11.7% to 20.8%) [47]. From 2011 to 2018, the usage rate among high schoolers increased nearly 14 times (1.5 % to 20.8%) [47]. Even among middle schoolers, usage rate increased over 8 times from 2011 to 2018 (0.6% to 4.9%) [47]. While the health effects of tobacco are well-known, much less is known about e-cig. In recent days, starting from late June, the surge in COVID-19 across the U.S. has been significantly attributed to increased incidence amongst the younger segment of the population. Given the prevalence of e-cig vaping in the young population, significant attention must be paid to how e-cig use may be related to COVID-19 incidence and severity.

Originally marketed as a safer alternative to traditional cigarettes, alarming health risks of e-cig have been slowly revealed. E-cig could be cytotoxic to both endothelial and epithelial cells and may decrease immune functions [48]. However, study of e-cig is complicated by the presence of nicotine and flavorings. Not all e-cig has nicotine and flavoring, but almost all brands have one or the other. These components can be significantly more cytotoxic than the base ingredients of e-cig, which is propylene glycol (PG) and vegetable glycerin (VG). While nicotine is known to be cytotoxic, we have previously reported that e-cig without nicotine can also induce DNA double strand breaks in cells [49].

In this study, we compared differences in the activation of key molecular pathways related to COVID-19 in tobacco and e-cig users. Specifically, we examined one cohort of e-cig users who only smoked e-cig without nicotine and without flavorings and another cohort of e-cig users who smoked e-cig with nicotine and with any flavorings. We found that tobacco use increases ACE2 expression, which corroborates the results of other studies [50]. However, e-cig use in any case did not increase ACE2 expression, which has not been reported in the literature. One recent study exposing mice to nicotine-containing e-cig for 1 month has found induction of ACE2 after exposure to nicotine-containing e-cig [51]. In our dataset, humans were also exposed to nicotine-containing e-cig for at least 1 month, suggesting that the in vivo results may not replicate physiological exposure.

We also found that tobacco use and use of nicotine/flavor-containing e-cig led to significant cytokine dysregulation and potential inflammasome activation. However, non-flavored and non-nicotine-containing e-cig use does not lead to either. While it has been reported that there is only limited inflammation in users of e-cig without nicotine and flavors [24], no study has compared gene expression changes of this group of e-cig users to users of nicotine/flavor-containing e-cig. One study also demonstrated inflammasome activation among e-cig users but did not take into account effects of nicotine and flavors [52]

Several limitations exist in our study that should be addressed in future studies. First, for the tobacco dataset, current smokers were only compared to former smokers, not never smokers. Second, for the e-cig dataset with flavor/nicotine-containing e-cig, only former smokers, not never smokers, were participants. However, since both the experimental and control group are composed of former smokers, the effect of smoking should not affect the statistical validity of our comparisons. Third, in the e-cig studies, participants were only exposed to e-cig for at least 1 month, while certain effects of e-cig may be expected to develop only after chronic exposure. Nonetheless, given that we observed cytokine/inflammasome dysregulations already given the relative short duration of exposure, we may expect chronic exposure to lead to even more dysregulation.

In conclusion, our study has demonstrated that tobacco and flavored or nicotine-containing e-cig use could both lead to increased inflammatory response, but only tobacco upregulates ACE2. Inflammation and ACE2 upregulation may increase susceptibility to COVID-19. While further experiments and epidemiology data are needed to confirm our results, we believe that our study is a critical early step into evaluating the implications of using tobacco and e-cig during the COVID-19 pandemic.

## 4. Materials and Methods

### 4.1. Datasets of Gene Expression from E-cig and tobacco samples

Bronchial epithelial cell sequencing data from e-cig and non e-cig users were obtained from GSE138326 (n=15 e-cig, 15 non e-cig), provided by Song et. al. [24], and GSE112073 (n=15 e-cig, 21 no e-cig), provide by Corbett et. al. [25]. In the Song et. al. dataset, 30 healthy subjects (21-30 years old) with no prior history of e-cig or tobacco use were randomly assigned to a e-cig vaping group or control group. In the e-cig vaping group, participants used the device twice a day at 20 puffs per session for exactly 4 weeks. E-cig was composed of 50% PG and 50% VG, with no nicotine or flavor. In the Corbett et. al. dataset, patients were aged 18-55 years old and were all former cigarette smokers, defined as having been tobacco-abstinent for over 3 months and have smoked at least 5 cigarettes for 2 years or more at some point of their lives. 15 subjects are former tobacco smokers who have used nicotine-containing e-cig of any brand or flavoring for at least 6 days a week and at least 1 month. 25 subjects are former tobacco smokers who have not used any nicotine-replacement device. Bronchoscopies were performed on all subjects for both studies, and bronchial brushings from the epithelium were profiled by microarray. RNA sequencing data for current and former tobacco smokers were obtained from adjacent normal samples of 49 lung squamous cell carcinoma (LUSC) patients from The Cancer Genome Atlas (TCGA).

### 4.2. Differential expression analysis

Differential expression was assessed between LUSC samples from current smokers and former smokers using the Kruskal-Wallis test (p<0.05). Microarray data from the e-cig datasets were analyzed using both the GEO2R software, which employs the limma (Linear Models for Microarray Analysis) R package, and the Kruskal-Wallis test (p<0.05). Non-smoking control vs. e-cig samples and samples from former smokers vs. former smokers who smoke e-cig were the comparisons made for the two studies.

### 4.3. Inference of immune cell infiltration populations using Cibersortx

The Cibersortx algorithm was be used to deconvolute microarray and RNA-sequencing data to estimate the infiltration levels of 22 immune cell types [26]. The immune cell types examined include, naïve B-cells, memory B-cells, plasma cells, CD8 T-cells, CD4 naïve T-cells, CD4 memory resting T-cells, CD4 memory activated T-cells, follicular helper T-cells, regulatory T-cells, gamma-delta T-cells, resting NK cells, activated NK cells, monocytes, M0-M2 macrophages, resting dendritic cells, activated dendritic cells, resting mast cells, activated mast cells, eosinophils, and neutrophils.

### 4.4. Correlation of smoking status/e-cig vaping status to pathways or signatures using GSEA

GSEA was used to determine the correlation to biological pathways or signatures for the same comparisons described in Section 4.1 [53]. A categorical phenotype file was used, with e-cig users, tobacco users, or control as labels. Gene sets were obtained from the Molecular Signatures Database (MSigDB) [54]. Canonical pathways (C2) and immunological signatures (C7) were correlated with vaping/smoking status (p<0.05).

### 4.5. Selection of cytokine and cytokine-related genes for analysis

Cytokines and cytokine-related included chemokines, chemokine receptors, interleukins, interleukin receptors, interferons, tumor necrosis factors (TNFs), and TGFB family members.

### 4.6. Selection of inflammasome-related genes for analysis

Both genes for proteins that constitute the inflammasome complex and genes that regulate the expression of these inflammasome components are included in our analyses. The former includes inflammasome sensor proteins, such as NLRP family proteins, AIM2, and NAIP; adaptor proteins, such as ASC (PYCARD); or effectors, including caspase proteins [40]. The latter includes NF-kB, IKK, MAPK3, and TAK1 [55].

## Author Contributions

Conceptualization, W.M.O.; methodology, W.M.O.; software, N/A; validation, A.C.L., J.C., and W.T.L; formal analysis, A.C.L., J.C., C.C. and W.T.L; investigation, A.C.L., J.C., C.C., and W.T.L; resources, W.M.O.; data curation, N/A; writing—original draft preparation, A.C.L., J.C., and W.T.L; writing—review and editing, A.C.L., J.C., W.T.L., E.Y.C., and J.W.R.; visualization, A.C.L., J.C., and W.T.L.; supervision, W.M.O.; project administration, W.M.O..; funding acquisition, W.M.O.. All authors have read and agreed to the published version of the manuscript

## Funding

This research was funded the University of California, Office of the President/Tobacco-Related Disease Research Program Emergency COVID-19 Research Seed Funding Grant (Grant number: R00RG2369) to W.M.O. and the University of California, San Diego Academic Senate Grant (Grant number: RG096651) to W.M.O.

## Acknowledgments

None.

## Conflicts of Interest

The authors declare no conflict of interest.

## Abbreviations

PG: propylene glycol
VG: vegetable glycerin
LUSC: lung squamous cell carcinoma
MSigDB: Molecular Signatures Database
TNF: tumor necrosis factor
limma: Linear Models for Microarray Analysis
Treg: Regulatory T cells
PBMCs: Peripheral Blood Mononuclear Cells
TCGA: The Cancer Genome Atlas

## Notes

### Competing Interest Statement

The authors have declared no competing interest.

